# Proton egress pathway during the S_1_–S_2_ transition of the Oxygen Evolving Complex of Photosystem II

**DOI:** 10.1101/2021.01.29.428861

**Authors:** Divya Kaur, Yingying Zhang, Krystle M. Reiss, Manoj Mandal, Gary W. Brudvig, Victor S. Batista, M. R. Gunner

## Abstract

Photosystem II uses water as the ultimate electron source of the photosynthetic electron transfer chain. Water is oxidized to dioxygen at the Oxygen Evolving Complex (OEC), a Mn_4_CaO_5_ inorganic core embedded in the lumenal side of PSII. Water-filled channels are thought to bring in substrate water molecules to the OEC, remove the substrate protons to the lumen, and may transport the product oxygen. Three water-filled channels, denoted large, narrow, and broad, that extend from the OEC towards the aqueous surface more than 15 Å away are seen. However, the actual mechanisms of water supply to the OEC, the removal of protons to the lumen and diffusion of oxygen away from the OEC have yet to be established. Here, we combine Molecular Dynamics (MD), Multi Conformation Continuum Electrostatics (MCCE) and Network Analysis to compare and contrast the three potential proton transfer paths during the S_1_ to S_2_ transition of the OEC. Hydrogen bond network analysis shows that the three channels are highly interconnected with similar energetics for hydronium as calculated for all paths near the OEC. The channels diverge as they approach the lumen, with the water chain in the broad channel better interconnected that in the narrow and large channels, where disruptions in the network are observed at about 10 Å from the OEC. In addition, the barrier for hydronium translocation is lower in the broad channel, suggesting that a proton from the OEC could access the paths near the OEC, and likely exit to the lumen via the broad channel, passing through PsbO.

## 1. Introduction

Photosynthetic organisms reduce atmospheric CO_2_ to sugar, generating oxygen as a byproduct, a process driven by solar light as the energy source and water as the ultimate source of reducing of equivalents (i.e., electrons). The water oxidation reaction is catalyzed by the oxygen evolving complex (OEC) of photosystem II (PSII), an inorganic Mn_4_CaO_5_ cluster ligated by the side chains of surrounding amino acid residues [1–4] (Figure 1C). The OEC cycles through five increasingly oxidized states, S_0_, S_1_, S_2_, S_3_ and S_4_ and forms the OO bond of O_2_ from two substrate water molecules during the transition from S_4_ to S_0_ [1–3,5,6]. The overall water oxidation reaction during the light period of photosynthesis in PSII is:

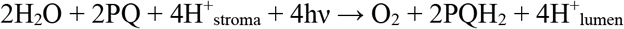

resulting in the release of four protons to the lumen while the protons needed for reduction of the two equivalents of plastoquinone (PQ) to plastoquinol (PQH_2_) are taken up from the stroma. Thus, the reaction generates a carrier of reducing equivalents (PQH_2_) and a transmembrane pH free energy gradient essential to drive the subsequent steps of photosynthesis [7]. In particular, PQH_2_ reduces the cytochrome b6f complex, which in turn passes electrons to flavodoxin via PSI and then to NADH at the end of the light activated photosynthetic electron transfer chain.

**Figure 1.**
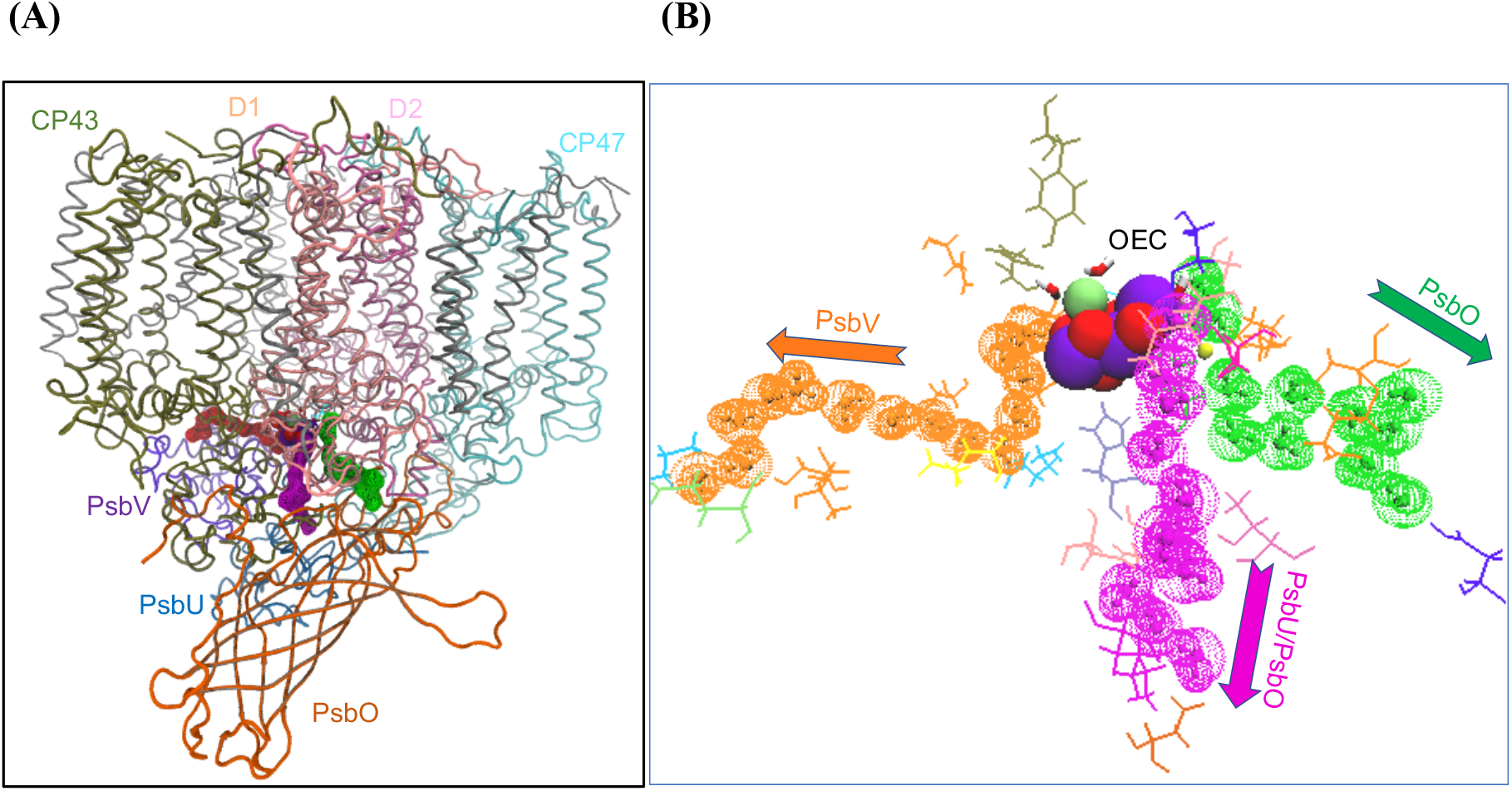
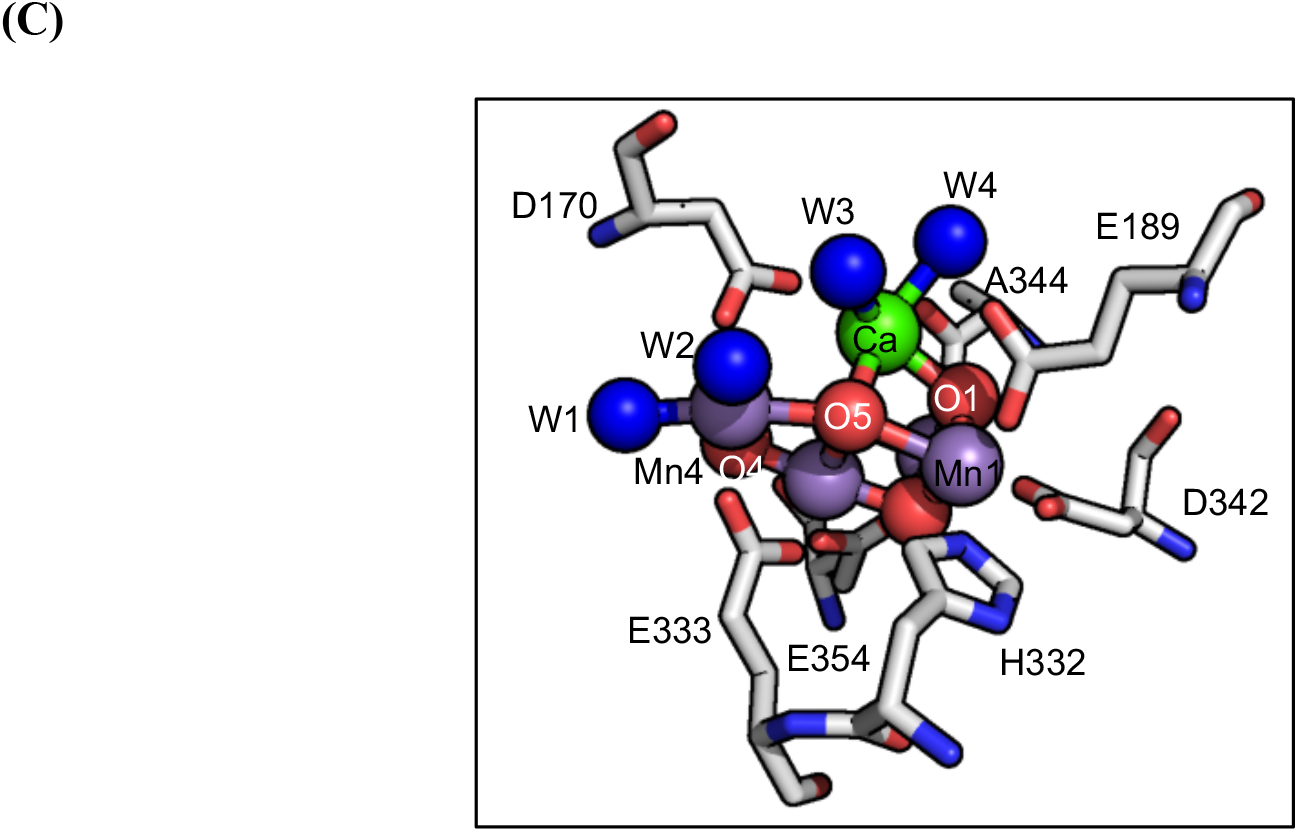
Structure of PSII and water channels. (A) Monomeric PSII highlighting D1, D2, CP43, CP47 and PsbO subunits (4UB6). The OEC cluster and water channels are shown as spheres. The full protein embedded in a lipid bilayer is included in the MD trajectory. (B) Water molecules in the large (orange), narrow (magenta) and broad (green) channels taken from one snapshot in the MD trajectory. (C) Crystal structure of OEC (PDB: 4UB6 [4,50]) with all terminal ligands. W1 and W2 are ligands to Mn4 and W3 and W4 to Ca. O5 is nominally closest to the broad channel entry, O4 to the narrow channel and O1 to the large channel.

The OEC is embedded in D1 protein subunit at about 20 Å from the lumen, the outer, positive side of the chloroplast. Thus, substrate water molecules must be delivered to the OEC and the products, including protons and O_2_, must be removed. However, the underlying mechanisms for water supply and product removal remain to be established. The crystal structures show long, complex water filled, hydrogen bond networks that might serve for the required functionality [4], including three channels commonly known as narrow, broad and large channels (Figure 1A and Figure 1B) [8–12].

Water diffusion from the lumen to the OEC likely requires water-filled channels. Diffusion of O_2_ away from the OEC could happen even through relatively hydrophobic paths [13,14], while protons typically diffuse through the Grotthuss mechanism [15].

Side chains are classified based on their possible roles in proton transfer. In Grotthuss proton transfer the proton defect is passed through highly-organized hydrogen bond networks [16]. Grotthuss proton transfer occurs when a proton is transferred to a lone pair on an adjacent water molecule or residue with which it was forming a hydrogen bond. An excess proton can thus move rapidly through a pre-aligned water chain without a hydronium ever being formally localized anywhere along the pathway. The Grotthuss-competent residues, serine, threonine and tyrosine, have a lone pair of electrons to accept a proton as well as a hydrogen-bonded proton to pass onward [17].

Ionized Asp or Glu residues have no dissociable proton, while an ionized His, Lys or Arg have no lone pair to accept a proton. However, they can serve as proton loading sites, groups that facilitate proton transfer by alternating between being “loaded” and “unloaded” intermediates, transiently gaining or losing a proton [7]. The acidic and basic residues facilitate proton transport by alternating between loaded/unloaded pronation states. Asn, Gln and Trp can make hydrogen bonds and anchor and orient the hydrogen bonded chain but are unlikely to serve as active intermediates in proton transport.

### Source of the released protons

In the oxidation of two waters to oxygen, four protons are lost to the lumen. The PSII XFEL structures provide insights into how the structure changes through the S-state cycle [6,18]. DFT-based QM/MM simulations have optimized the OEC structure, hinting at the sites of proton binding within the cluster through the reaction cycle [19]. The source of the lost proton will be different in each S-state transition. Going from S_0_ to S_1_, the donor is likely to be a bridging oxygen, releasing a proton retained by the OEC following water oxidation in the S_4_-state of the previous Kok cycle. Thus, its location may provide information about the position of the substrate waters bound to the OEC prior to O_2_ release [1,19,20]. EPR [21] and associated computational studies [22–24] favor a protonated O5 bridge. However, other studies have supported a protonated O4 [25] or O1 [26]. In all other S-states, which are more oxidized than S_0_, none of the cluster-bridging oxygens retain a proton. There is little or no proton release to the lumen when S_1_ is oxidized [27–29]. In the oxidation states beyond S_2_, protons have been proposed to be released from the terminal waters of Mn4 (W1, W2) or of Ca (W3, W4) or from waters, such as WX bound nearby. The resulting hydroxyl may inserted into the cluster as a substrate for O_2_ formation [3,24,30–33].

The question remains what role each channel plays in the transport of the substrate and products [3,9,12,34] and each has been considered as a proton exit pathway. The broad channel nominally originates from the O5 bridge in the vicinity of Mn4 and moves to the lumen via the peripheral PsbO subunit (Figure 1). Continuum electrostatic simulations [10] showed a monotonic increase in proton affinity of acidic and basic residues moving along the broad channel to the lumen, suggesting this channel could provide a downhill path for proton exit. MD studies [34–36] of how the PsbO surface is connected to the broad channel waters also shows that acidic residues in this channel can serve as proton loading sites facilitating proton transport. Recent, eigenvector centrality analysis using a MD trajectory [37] on a 25-Å sphere centered at the OEC in the S_1_ and S_2_ states find that the narrow and broad channel are favorable for water and proton transport while the large channel may be better suited for transfer of larger ions.

The narrow channel has been assigned as originating from the OEC O4 bridge and includes the terminal water ligand, W1, of Mn4. It travels to the lumen alongside the broad channel through the interface of the PsbU and PsbO subunits (Figure 1A and Figure 1B). Several computational studies of the S_0_ to S_1_ transition [9,25] favored release of a proton from O4 which appears to exit towards this channel. The oriented waters entering the narrow channel make it look well-prepared to transfer protons via a Grotthuss mechanism [25,37]. However, small, neutral, polar molecules such as ammonia [38–43] and methanol [44,45] bind in this channel, suggesting that it could be the path for substrate water delivery. Steered MD studies [11] have also shown the narrow channel has the lowest barrier to water permeation. If W1 is a substrate water, it would then be well positioned to be replenished via this channel.

The large channel originates from O1, reaching the lumen in the PsbV subunit (Figure 1A and Figure 1B) [11]. It contains more hydrophobic residues than the other two, so has been proposed as the path for oxygen release [11]. MD studies [9,11] show that this channel has the highest barrier to water permeation. However, O_2_ need not transfer through a water-filled channel and solvent accessible cavity analysis [9] and noble gas derivatization [8] proposed a ‘back channel’ as a separate, more hydrophobic pathway for the O_2_ release. Experimental [20,30] and computational studies [11] have also favored the large channel for substrate water delivery. The large channel appears to be the closest point of entry if terminal water ligands to the OEC Ca serve as a substrate [20,30,46,47]. This channel has also been proposed to play a role in proton transfer [34,48].

Thus, the current view of the region around the OEC sees three distinct channels, each of which has been proposed to play a role in water delivery and proton release. While, a consensus may be forming that the broad channel is the proton exit channel [10,34] and the narrow channel carries water [11] and the large channel oxygen [11], different experiments and simulations have assigned different roles to each. Thus, it may be that any of the channels can be bifunctional or that the substrates and products can use multiple paths moving between the OEC and lumen [10,25,49].

Here we compare ability of the three water-filled channels to transport a proton upon formation of the S_1_ state [23,50]. The connection between the three channels near the OEC and the barrier to hydronium transfer through the channels are studied by combined Molecular Dynamics (MD), Multi Conformation Continuum Electrostatics (MCCE) [51] and Network Analysis [52]. The main conclusions are that all three water channels are highly interconnected near the OEC. In addition, the energy to place a hydronium in any putative water channel is also very similar near the OEC. However, moving out towards the lumen, the broad channel is energetically more hospitable for a hydronium than the large and narrow channel. Thus, a proton can exit from any point on the OEC to leave the protein via the broad channel. Likewise, the interconnections of the water channels near the OEC suggest substrate waters can enter via any channel to provide the substrate to any point on the OEC.

## 2. Methods

### 2.1. MD protocol for system preparation for wild-type PSII

The MD simulations start with the 1.9 Å resolution X-ray crystal structure of *Thermosynechococcus vulcanus* 4UB6 [4]. The 20 subunits of one PSII monomer are included. The CHARMM-GUI [53] bilayer membrane builder embeds the system within a membrane of MGDG, DGDG and POPG in 1:1:1 ratio. The final system has the PSII monomer in a 180 Å × 180 Å × 81 Å rectangular box with ~300,000 TIP3P [54] water molecules along with the membrane. There are 616 Na^+^ and 320 Cl^−^ ions added to maintain charge neutrality at a 0.15 M concentration. There are 494,990 atoms in the system.

The OEC bond lengths, angles and dihedrals are taken from the QM/MM optimized S_1_ state [50]. The structure is maintained with constraints of 1000 kcal/mol/Å^2^ for bond lengths and 200 kcal/mol/Å^2^ for angles. Additional restraints of 500 kcal/mol/Å^2^ are added for the hydrogen bonds between Q_B_ and Ser264 and between Y_Z_ and H190 [55]. The latter may be a strong, short hydrogen bond, which will not be maintained in the standard classical mechanics force field [56]. ESP charges are used for the Ca^2+^ and four Mn (Mn1, Mn2, Mn3 and Mn4). RESP charges are used for the 5 μ-oxo ligands, four terminal waters of Ca^2+^ and Mn4 as well as the primary OEC ligands D1-D170, E189, H332, E333, D342, A344, and CP43-E354. The ESP charges [37] are obtained using density functional theory at B3LYP/6-31G level using the Gaussian software package [57]. RESP charges were derived from the DFT-calculated ESP charges using the RESP program [58]. CHARMM forcefield parameters for lipids and for neutral pheophytin, chlorophyll and plastoquinone cofactors of PSII were taken from Guerra et al. [56].

Patching is carried out for both heme b_559_ and the non-heme Fe complex in CHARMM [59]. The di-sulphide linkage [60] between Cys 19 and 44 in PsbO is made. The protonation states of the residues must be fixed at the beginning of the MD simulations. The ionization states of acidic and basic residues were modified based on MCCE calculations of the protein in the S_1_ state with all other cofactors in their ground state [27]. All Asp, Glu, Arg, and Lys are ionized and His, Cys and Tyr are neutral with the following exceptions: CP47 -D380, E387, E405, CP43 -E413, D2 -E242, E343, PsbO- D102, D224, E97, PsbV- K47, K134 are neutral, while His D1- H92, H304, H337, CP47- H343, CP43- H74, H398, PsbO- H228 and PsbU-H81 are protonated. In addition, by default CHARMM-GUI chooses the neutral His with a proton on ND1 (HSD). However, MCCE found that the following His residues prefer to have a proton on NE2 (HSE): D1-H195, H252, CP43- H157, H201, D2- H61, H87, H189, H336, PsbO- H231, PsbV- H118. It should be noted that these choices have consequences. For example, choosing the HSD tautomer for D1-His252 leads to the opening of a loop near Q_B_.

OpenMM [61] is used to generate MD trajectories including periodic boundary conditions with Langevin dynamics with Nose-Hoover Langevin piston at constant pressure (1 bar) and temperature (303.15 K). The coulombic interactions are calculated using the Particle-Mesh Ewald (PME) algorithm. A 2fs time step is used. To relax initials clashes, the system is first minimized in CHARMM using Steepest Descent (SD) minimization for 1000 steps. The system is then further equilibrated in OpenMM where the protein backbone, side chains and lipids are allowed to relax for 355 ps when all constraints are removed. The production run is 100 ns long.

### 2.2. MCCE calculations

Multi-Conformation Continuum Electrostatics (MCCE) [51] calculations are carried out on 10 snapshots from the 100 ns S1-state MD trajectory. The calculations are focused on the 88 Å × 70 Å × 65 Å rectangular region around the OEC and the regions of the protein that connect the OEC to the lumen (Figure 1A). These snapshots are chosen with the MDAnalysis clustering algorithm [62] and represent structural clusters based on the distances between D1-D61, E65, D2-E312, K317, PsbO-D224 and Cl^−^.

MCCE [51] uses the continuum electrostatics program Delphi to calculate electrostatic energies from the Poisson-Boltzmann equation. The parameters and methods have been described previously [26,27]. The implicit salt concentration is 0.15 M with a 2 Å stern layer, and a pH of 6. The dielectric constant for the protein is 4 and dielectric constant for the solvent is 80. Parse charges [63], optimized for continuum calculations, are used for amino acids while integer valence charges are used for the OEC [27]. TIPS [54] charges are used for explicit waters. Standard MCCE topology files are used for the cofactor chlorophylls, pheophytin, plastoquinone, chloride, heme and non-heme iron. Amber van der Waal parameters are reduced to 25 % of their full value based on previous MCCE benchmark calculations [64]. As we are concerned with the proton exit from the OEC which is out of the intra-membrane region no lipids are included.

MCCE is always limited to a fixed backbone. Here we use iso-steric sampling, which finds the many hydrogen bond/ionization states in a Boltzmann distribution consistent with the heavy atom positions found in the input MD snapshot. Thus, the neutral and ionized protonation states of the Asp, Glu, Arg, Lys, His and Tyr residues are sampled. The amide terminal of Asn and Gln and neutral His tautomers can exchange. Hydroxyls of Ser, Thr and Tyr can reorient. Both chlorides, one near D2-K317 and other near D1-N338 and F339, are included in the calculations. The primary ligands of OEC, D1-E189, D1-E333, D1-D342 and CP43-E354, are fixed in their ionized state while D1-H332 is neutral [27].

The MCCE calculation retains waters within the MD snapshots with <15% solvent accessible area. All waters surrounding the protein are replaced by implicit solvent. The positions of the OEC atoms are retained along with the oxygen positions of the terminal waters ligated to Mn4 (W1 and W2) and Ca^2+^ (W3 and W4) of OEC. The positions of water oxygens within the protein are taken from the input MD snapshot. Each water oxygen can choose between 15-20 different proton positions in Monte Carlo (MC) sampling and can sample neutral H_2_O and hydronium H_3_O^+^ conformers. Except for the terminal waters of the OEC, all waters can be moved into solution. We find in the average of all accepted microstates 92% of these waters remain bound in the protein, with few having no occupancy.

#### 2.2.1 Hydrogen bond network analysis

MCCE hydrogen bond analysis focuses on the 88 Å × 70 Å × 65 Å rectangular region around the OEC including the regions of the protein that connect the OEC to the lumen (Figure 1A). For each snapshot, the calculations sample ≈33 million microstates during MC sampling, giving the distribution of protonation states, water occupancy and polar hydrogen positions. These are analyzed to obtain the hydrogen bond connections between the residues. Figure S1 shows the region of the protein analyzed within the whole PSII.

The combined MCCE-network hydrogen bond analysis has previously been used to analyze proton transfer pathways in cytochrome c oxidase and complex I, and the same methods are used here [13,65]. Hydrogen bonds have a distance between the donor, hydrogen (D-H) and acceptor (A) of 1.2 Å - 3.2 Å and an angle between D-H and A > 90°. The connection is counted if the proton acceptor and donor form a hydrogen bond in > 0.1 % of accepted microstates in the MCCE simulation starting with a given MD snapshot. Cytoscape [52] draws the network (Figure 2C). Residues are nodes and each line indicates the formation of a hydrogen bond, either directly connecting two residues or via as many as four bridging waters. Longer water chains do not find additional connected residues. Residues connected by four waters can be as much at 10-13Å apart.

**Figure 2.**
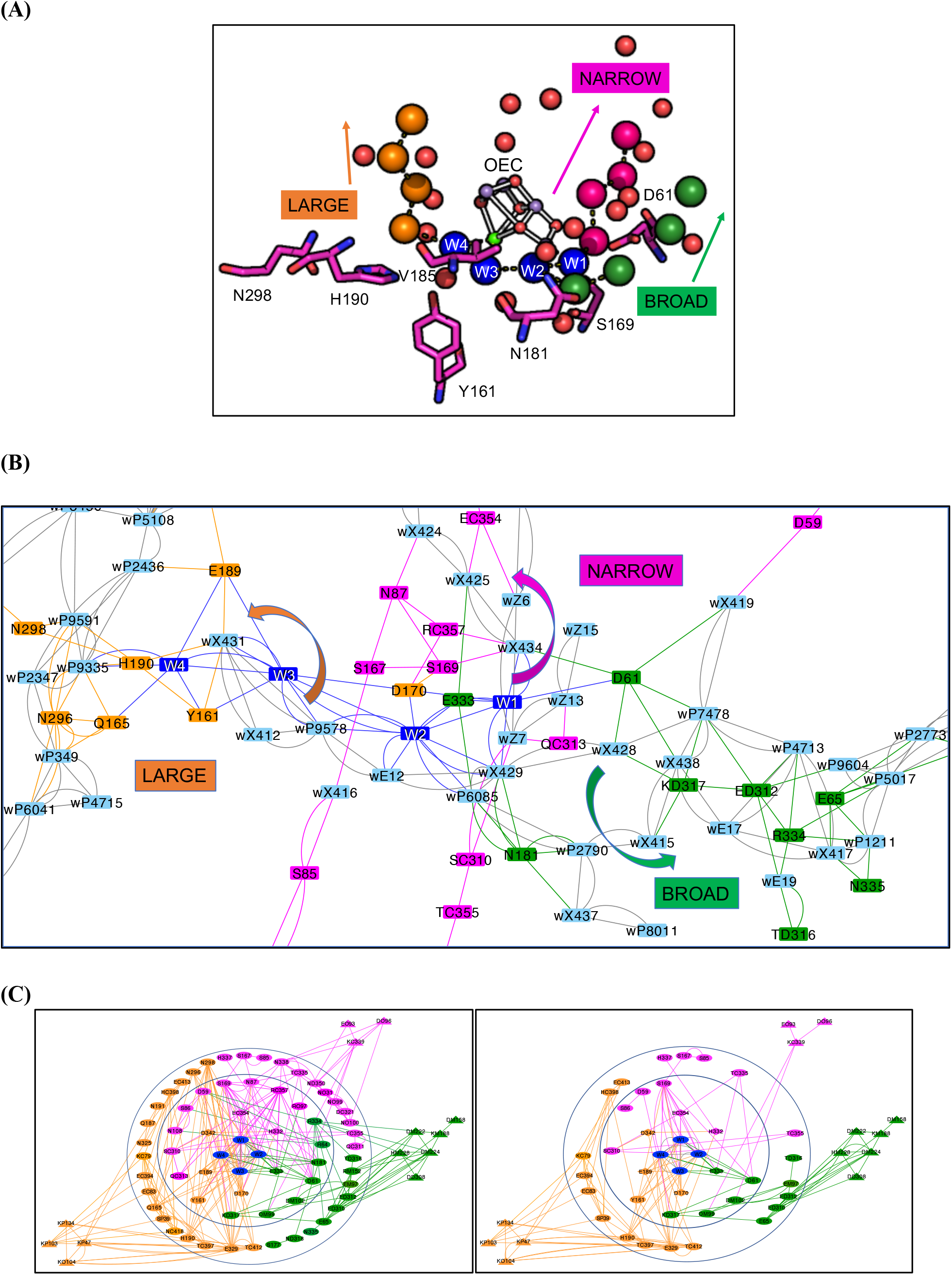
Three views of the hydrogen bond network surrounding the OEC. The colors are consistent between all figures. Orange: large channel; green: broad channel; magenta: narrow channel. Dark Blue: Primary water ligands to Mn4 (W1, W2) and to Ca (W3 and W4). (**A)** Water oxygen atoms and hydrogen bonded residues within 7Å of the OEC. The arrows indicate the direction of each channel. Water oxygens are colored by the nearest channel. Red spheres are waters that are not assigned. Residues associated with the Narrow channel: D1-S169; Broad channel: D1-D61, N181, V185; Large channel: D1-Y161, H190, N298. (**B**) The highly interconnected region near the OEC from the MCCE/network analysis for one representative MD snapshot. Water molecules are light blue and are labeled w. Water molecules X, E and Z are derived from the crystal structure while those labeled P are added during MD set up with an arbitrary number. The coordinates for the input snapshot can be found in the SI. Hydrogen bond connections made in > 1% of the MC accepted states are shown by connecting lines. Arrows show direction of proton transfer towards the three separated channels. Table 2 lists several possible paths though the network to the lumen from each of the four OEC terminal waters. (**C**) Network of hydrogen bond connections from the OEC to the lumen found in MCCE calculations starting from a representative MD snapshot. Nodes are labelled as: Residue type, Chain designation, Residue number. No chain designation indicates D1. Chain C is CP43; D is D2; M is PsbO; O is PsbU; and P is PsbV. For example, EC354 is CP43-Glu354. Diamonds are primary ligands and triangles are residues with at least 20% of their surface exposed to the lumen. Lines show hydrogen bond connection mediated by four or fewer waters. The inner circle encloses highly interconnected residues near the OEC. Connections between residues nominally in different channels are seen. The outer circle encloses residues in their separated channels. Beyond the outer circle are residues connecting the channel and the surface. *Left*: Nodes that contain all residues in Table 1. *Right*: Nodes that show only residues that can participate in Grotthuss proton transport or serve as proton loading sites, labeled PT in Table 2. ^§^Figure S2 provides network figures for nine additional snapshots for comparison.

**Table 1.** The residues found in the hydrogen bond network; Residues are divided by their properties: PT are capable of direct participation in proton transport by Grotthuss proton transfer or by serving a proton loading site by being transiently protonated/deprotonated; Polar, non-Grotthuss: can make hydrogen bonds to anchor the network but are unlikely to carry a proton towards the lumen; Non-polar: residues associated with the channels that have been the subject of prior investigation. Residues are also separated by their location: Origin: region of the OEC that is closest to the channel entrance; Interconnected near OEC: region including residues shown in Figure 2B; Separated Channel: residues committed to an individual channel; Surface residues: residues with surface >20% surface exposure. *Primary ligands of the OEC. ^#^surface residues connected to multiple channels. The colors assigned to each channel are used in all figures. Recently the narrow, broad and large channels were referred to as O4, O5 and O1 channels, indicating the atoms in the OEC where they are proposed to originate [18].

| Channel | Type | Origin | Interconnected near OEC | Separated channel | Surface residues |
| --- | --- | --- | --- | --- | --- |
| Narrow Magenta | PT | *O4, Mn4-W1* | D1-D59, S86, S169 [68], CP43- S310, H332*, E354* | D1-S85, S167 (near OEC), D1- H337, CP43- T335, T355, D321 | PsbU-E93, D96, CP43-K339 |
|  | Polar non-Grotthuss |  | D1-N87 [69], N108, CP43-Q313, R357, PsbU-R97 | D1- N338, CP43-Q311, PsbU-N31, N99, N100, D2-N350 |  |
|  | Non-polar |  | D1-P334 |  |  |
| Broad Green | PT | *O5, Mn4-W1, W2* | D1-D61 [70], D2-K317 [27,71], PsbO-D99, D102, E333* | D1-S177, E65 [72], D2-E310, E312, D2- Y315<sup>§</sup>, T316, PsbO-E97, D99, D102 [34] | D2-D308, PsbO-D158, K188, D222, D223<sup>§</sup>, D224, H228, E229<sup>§</sup> |
|  | Polar non-Grotthuss |  | D1-R64, N181, R334 | D1-N335, D2-N318, PsbO-R152 |  |
|  | Non-polar |  | D1-V185 [73] |  |  |
| Large Orange | PT | *O1, Ca-W3, W4* | D1-Y161, D170*, E189*, D342* | D1- H190, E329, CP43-E83, E413, E394, T412, H398, K79, T397, PsbV-S39 | PsbU-K47, K134, PsbV-K103, K104 |
|  | Polar non-Grotthuss |  |  | D1-Q165, Q187, N191, N296, N298 [48,74], N325, D2-R348<sup>§</sup>, CP43-N418 |  |
|  | Non-polar |  | PsbV-L341 |  |  |

**Table 2.** Example paths from each of the four OEC waters to each of the three channels.

|  | W1 | W2 | W3 | W4 |
| --- | --- | --- | --- | --- |
| Narrow | X434, | X429, W1, X434, | P9578, E12, X429, W1, X434, | W3, W2, W1, X434, |
|  | *All paths: X425, X424, Z4, P6395, Z21, P5192, P4445, P5284, P431, P5049, <u>K339</u>* |  |  |  |
| Broad | X429, | X429, | P9578, E12, X429, | W3, P9578, E12, X429, |
|  | *All paths: X428, P7478, P4713, X417, P1211, P5017, P9096, P8101, <u>D224</u>*<br>*Alternate path1: X428, <u>D61</u>, P7478, P4713, <u>E312</u>, P9604, P2773, P1510, P9096, <u>D224</u>*<br>*Alternate path2: X428, <u>D61</u>, P7478, P4713, X524, <u>E65</u>, P2577, P8298, P6850, <u>D224</u>* |  |  |  |
| Large | X429, E12, P9578, W3, W4, P9335 | X429, E12, P9578, W3, W4, P9335 | W4, P9335, | P9335, |
|  | *All paths: P9335, **P2436**, P5108, P2656, P9584, P815, P6926, P8089, P3306, P6393, P9357, P4873, X447, P7214, P4347, P8277, <u>K103</u>*<br>*Alternate path1: **P2436**, P5108, <u>E329</u>, P815, P6926, P8089, P3306, P6393, P9357, P4873, X447, P7214, P4347, P8277, <u>K103</u>*<br>*Alternate path2: **P2436**, P5108, <u>E329</u>, P815, P6926, P8089, <u>E413</u>, P4677, P4873, X447, P7214, P4347, P8277, <u>K103</u>* |  |  |  |

#### 2.2.2 Estimate of the energy of hydronium transferring through the channels

The free energy of hydronium is calculated at pH 6 in the three channels. This calculation may be viewed as the Monte Carlo analog of a PMF calculation in MD [66]. The OEC is fixed in the S_1_ state where Mn1 and Mn4 are in the +3 oxidation state while Mn2 and Mn3 are +4. One water at a time is fixed to be hydronium and the position (but not the charge) of nearby residues and water are allowed to come to equilibrium. In the MCCE calculations, all Asp, Glu, Arg and Lys are ionized and His are neutral except residues: D1-H92, H337, CP43-H91, H398, PsbO-H228, and PsbU-H81. The reference energy adds the energy of hydronium in solution (13.2 kcal/mol at pH 6) to the total MCCE energy of PSII relaxed without the hydronium. This is compared with the PSII energy with the hydronium in each position walking from the OEC to the lumen. Recent benchmark calculations [66] find good agreement for the energy barrier for proton transfer in the gramicidin channel with MCCE simulations and that previously found using the Empirical Valence Bond (EVB) model [67]. The calculations of hydronium are done on 8±2 positions of water in each snapshot.

## 3. Results

### 3.1. Analyzing the hydrogen bond network near the OEC

The OEC is ~20 Å from the lumen. In the work presented here the connections from the OEC to the surface are traced by combined MD/MC/Network Analysis. MD trajectories allow the protein to explore conformational space, while the MC analysis provides the Boltzmann distribution of hydrogen bonds (i.e. those that are energetically accessible), as well as residue protonation states, that are consistent with the heavy atom positions in individual MD snapshots. Networks include all internal waters from the MD snapshot. As will be shown, the network is dominated by water molecules, but amino acid side chains play significant roles defining the water-filled channels.

### 3.2. Representations of the pathways from the OEC to the lumen

Figure 2 shows the hydrogen bond network around the OEC. Residues and waters are colored by their assignment to a given channel (Table 1). Figure 2A shows a conventional view of the polar residues and waters within 10±2 Å of the OEC. This highlights the separation of the channels going away from the OEC. However, Figure 2B shows the multiplicity of connections, largely through waters, within 10±2 Å of the OEC, while Figure 2C provides a more schematic view, with only the amino acids that are hydrogen bonded via waters reaching from the OEC to the lumen. Each figure highlights the complexity of the possible pathway for a proton in different ways.

### 3.3. Protons released from any site on the OEC can enter any channel via interconnected waters

Figure 2B shows the results of the network analysis of the hydrogen bonds found by MC sampling of the polar proton and water positions in a single MD snapshot within 15 Å of the OEC. The channels are well separated beyond 10-12 Å of the OEC and these will be described below. However, near the OEC there are numerous connections amongst waters anchored by residues. Thus, here the network is better described as a river delta than as three separate channels. Table 2 provides examples of several paths from each primary OEC water ligand to the lumen. Nominally W1 is identified with the narrow and broad channels, W2 with the broad channel, and W3 and W4 with the large channel. However, a path can be found allowing each of the four terminal water ligands to exit via any of the three channels. The number of steps to reach a region where the channels are separated ranges from zero for W1 and W2 entering the broad channel or W4 to the large channel to five steps needed for W1 or W2 to enter the large channel.

### 3.4. The connections from the bridging oxygens in the OEC cluster

A proton is likely to be lost from one of the bridging oxygen in the S_0_ to S_1_ transition [3,23,25]. Figure 3 shows that with modest remodeling of the hydrogen bond network a proton on O1, O4 or O5 can enter into the water-filled channels. However, the primary ligands create a boundary around O2 and O3, with C-terminus D1-A344 blocking O2 and D1-H337 blocking O3, so they are poorly connected to the network.

**Figure 3.**
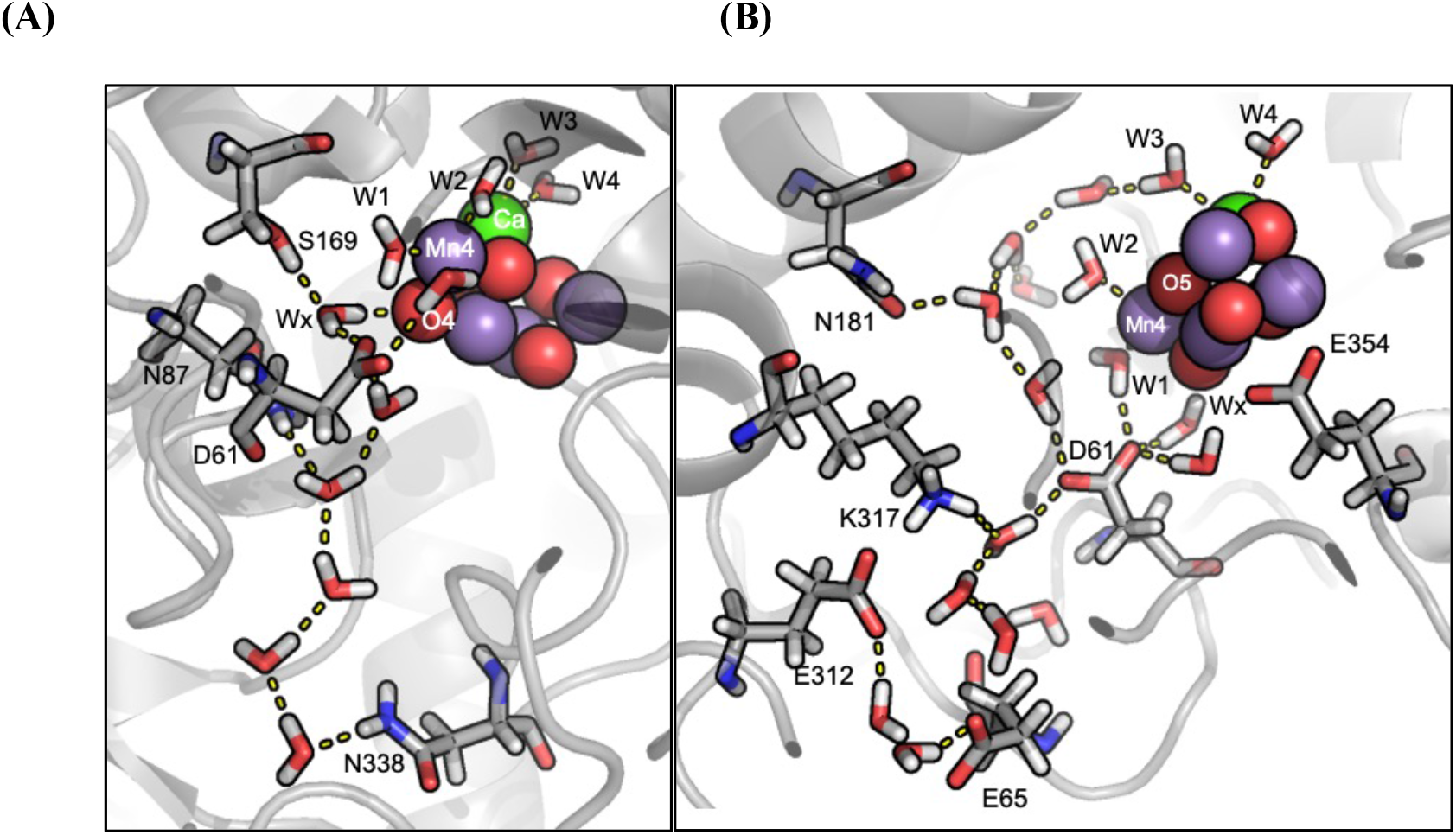
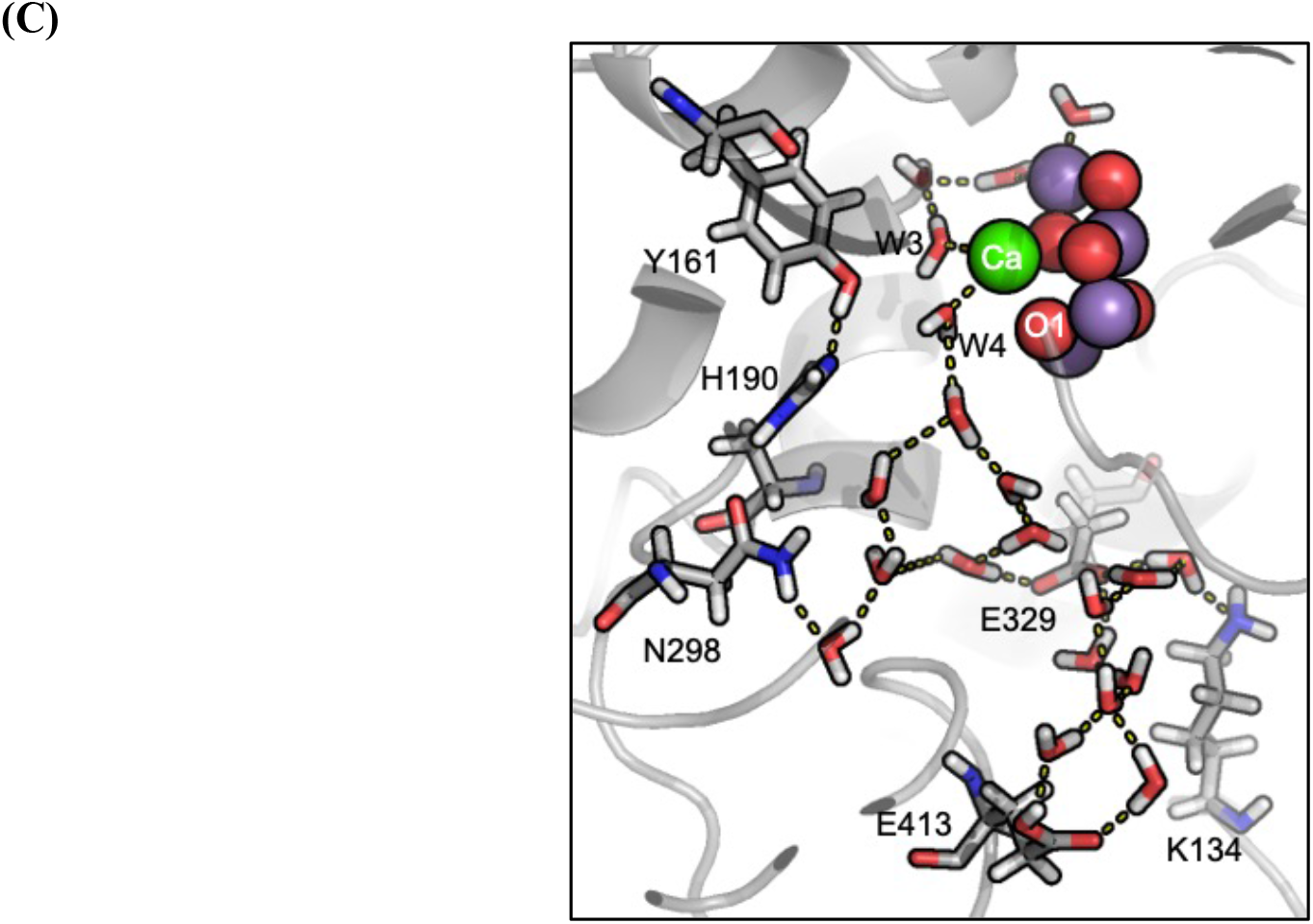
Representative hydrogen bonding pathways via ordered waters in a single MD snapshot. (A) Near O4 leading towards the narrow channel showing D1-D61, N87, S169, N338 (B) Near O5 leading towards the broad channel showing D1-D61, E65, N181, D2-E312, K317 and CP43-E354; (C) Near O1 leading towards the large channel showing D1-Y161, H190, N298, E329, CP43-E413 and PsbV-K134.

### 3.5. Amino acid participation in the network near the OEC

The connections available for proton transfer shown in Figure 2B and Table 2 are dominated by waters. Thus, Table 2 shows that there are paths to the lumen solely via waters without protons needing to hop though any amino acids. However, representative paths can be drawn with protons transferring via protonation/deprotonation of D1-D61 and D2-E312 to the broad channel or via D1-E329 and CP43-E413 to the large channel.

Figure 2C highlights the amino acid side chains making hydrogen bonds to waters within the channels. The network shown extends to the lumen. In the schematic network, a line connecting two residues can indicate anywhere from 0 to 4 intervening water molecules. The inner circle consists of the four water ligands, W1, W2, W3 and W4 that are in the center of the network. The four OEC water terminal ligands are surrounded by the six primary ligands. All primary ligands to the OEC Mn are connected to at least 6 other residues, sometimes directly but more often through complex water chains, with the exception of the Mn1 ligand D1-H332 which is isolated and only connected to H337. The next circle shows the residues near the OEC. While these are colored to indicate the channel with which they have been associated, they are interconnected by the complex water network shown in Figure 2B. The outer circle shows the residues in the well separated channels and the surface residues (triangles) where protons can exit. The peripheral residues connected to the outer circles make few connections to other residues in the network and are often on the surface. Networks obtained with different snapshots are provided in Supplementary Material (SI). They are drawn with the same layout so they can be easily compared. Peripheral residues that are unconnected in the network shown in Figure 2B are connected in the analysis starting with different MD snapshots. Overall, the network shows a total of 74 residues, with 23 residues identified in the broad channel, 26 residues in the narrow channel and 25 large channel residues (Table 1, Figure 2C).

Figure 2B and Figure 2C show the residues identified with each channel are linked near the OEC by multiple of possible connections. Different types of residues with different roles of proton transfer are found near the entry to the different channels. Thus, the narrow channel has more hydroxyl, Grotthuss competent residues and only one acid; In contrast, the broad channel has more potentially proton loading Asp and Glu as does the large channel. Each channel is identified with multiple Asn. The Asn’s amide side chain can anchor the network. We will give several examples that highlight the interconnections of residues near the OEC that have been shown to effect oxygen evolution.

### 3.6. The connections via D61

D1-D61, which bridges the narrow and broad channels, is a direct hydrogen bond acceptor of W1 and W_X_. D61 can help trap the proton being released from W1 as Mn4 is oxidized in the S_2_ state [27]. In different conformations in the Boltzmann ensemble, W_X_ makes hydrogen bonds to the O4 μ-oxo bridge as well as to D1-S169 and the D61 carboxylate. These waters connect D61 to residues identified with the narrow channel residues D1-S169, N87, N335, CP43-R357 (Figure 4A) and broad channel residues D1-N181, D1-R334, D2-K317 (Figure 4B) as well as all four OEC water ligands. Thus, a large number of amino acids can be connected to D61 either directly or via waters without any intervening amino acids.

**Figure 4.**
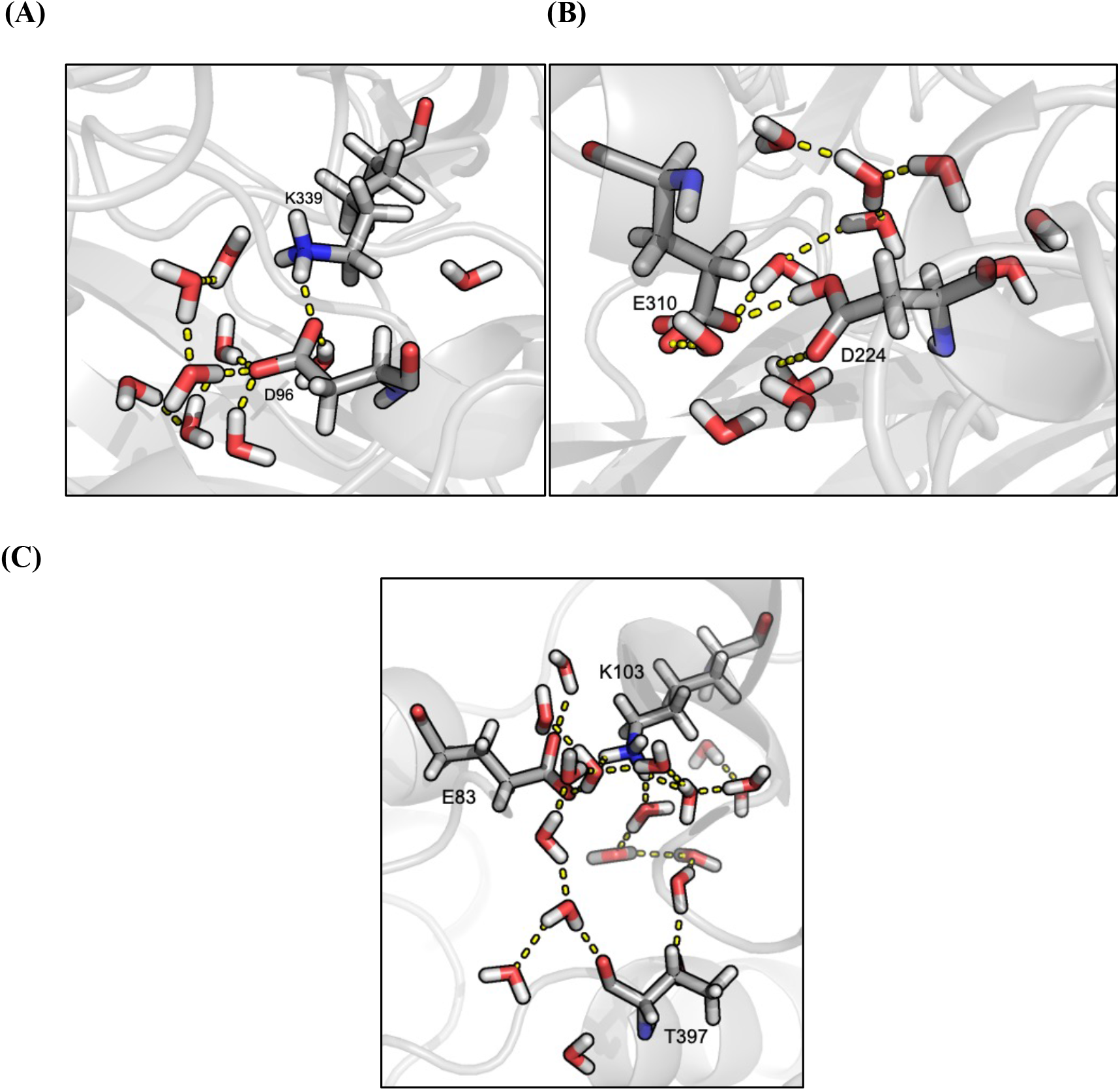
Network of hydrogen bonds from a single MD snapshot showing pathways through separated channels. (A) Narrow channel showing CP43-K339 and PsbU-D96. (B) Broad channel showing D2-E310 and PsbO-D224. (C) Large channel showing CP43-E83, T397 and PsbV-K103.

The D1-D61A mutation blocks the OEC from advancing beyond S_2_. The mutation modifies the FTIR spectra from the S_1_ to S_2_ and S_2_ to S_3_ transitions, supporting a role for this residue in the active proton transport network [70]. FTIR studies [75] find a large hydrogen bond network that includes D1-D61. For example, mutation of D1-E329, identified with the large channel, shows changes in the FTIR spectra similar to those associated with D1-D61 or with E65 and D2-E312 in the broad channel.

### 3.7. The connections near Y161(Y_Z_)/H190

Y_Z_ is the electron donor to P_680_ and it in turn oxidizes the OEC. On oxidation it transfers its hydroxyl proton to H190. The hydrogen bond between them is strengthened in the MD trajectory and is retained in the network analysis. The few waters around the YZ/H190 pair are not well connected to the larger hydrogen bond network surrounding the OEC. The nearby N296 and N298 are not Grotthuss conducting residues. Thus, the network analysis suggests poor egress opportunities for a proton bound between Y_Z_/H190. This is advantageous as the proton must not be lost from the His as it returns to the Tyr when the electron is transferred from the OEC.

### 3.8. The connections via D2-K317

D2-K317, near D61, and the Cl^−^ bound near the OEC are associated with the broad channel (Figure 3B). The Lys is connected to D61 via water molecules, and with D1-E65, N181, D2-E312 (broad channel) and with D1-D59 (narrow channel) as well as with the primary ligands D1-E333, D170 and E189 and all four OEC water ligands. D2-K317A or K317R mutants show reduced O_2_ evolution blocking/slowing advancement beyond S_2_ [71,75]. Simulations have shown that in the absence of Cl^−^, D61 and K317 form a salt bridge that can cut off the hydrogen bond network entering the narrow channel [76]. This further confirms the observation made in the D61A mutant, that being that the aspartate must be available the accept a proton in order for the OEC to proceed beyond S_2_.

### 3.9. The connections via D1-N181 and N87

D1-N181 is identified with the broad channel. It forms hydrogen bonds via waters to the primary ligands D1-D170, E189, E333 and CP43-E354 as well as to W2 of Mn4. The network analysis finds water mediated connection to the D1-D61, D2-E312, K317 (broad channel), to D1-D59, S169, CP43-R357 (narrow channel) and to D1-Q165 (large channel). An Asn is unlikely to directly participate in proton transport but can play a role in anchoring the hydrogen bond network. With the rotation of the terminal amide, the side chain can reorient the direction of its hydrogen bond donor and acceptor ends. MCCE finds this Asn to be oriented with the NH2, proton donating side oriented towards D1-D61. FTIR studies on N181A mutated PSII [75,77] found that this residue is part of the hydrogen bond network that includes D1-D61 described above.

D1-N87, which is identified with the narrow channel, forms water mediated hydrogen bonds to D1-S167, S169, N338, CP43-T335 and R357 (narrow channel), with D1-D61 (broad channel) and with the primary OEC ligands D1-E333, D170, CP43-E354 and W1 and W2. FTIR difference spectra [69] of D1-N87A mutated PSII look like those found for the wild-type PSII, although flash-induced O_2_ evolution studies show a decrease in the cycling efficiency.

### 3.10. The connections via CP43-R357 and D1-R334

CP43-R357 is associated with the narrow channel. The guanidinium group can donate as many as five hydrogen bonds to widely spaced neighboring waters or residues. The residue is hydrogen bonded via waters to D1-N87 (narrow channel), D1-D61 and N181(broad channel) and D1-Y161 (large channel), as well as with the primary ligands D1-E333, D170, E189, CP43-E354 and with the OEC water ligands. FTIR [75] studies find that R357 participates in a hydrogen bond network with the Ca and Mn water ligands. Previously, it was proposed [78] that this residue may be involved in proton transfer to the lumen beyond the S_2_ state. However, Arg has a very high intrinsic solution pK_a_ so is unlikely to become deprotonated in the protein [79].

R334 is associated with the broad channel. The network analysis finds D1-R334 hydrogen bonded to residues D1-D59, D61, E65, D2-E310, E312 and T316 (broad channel). Mutation of this residue finds changes in the efficiency of the S-state transitions beyond S_1_. FTIR spectra [80] on D1-R334A PSII shows the elimination of C=O peaks in S_2_-S_1_ and S_3_-S_2_ difference spectra. Also, the mutation decreases efficiency of the S_3_ to S_0_ transition.

In the inner region the pathways can be found through Figure 2B. Chain X, E and Z are crystal waters and chain P are waters added during MD set up. Amino acids on the paths are underlined. Waters that are common to all paths along one channel are in italics.

## 4. Hydrogen bonding network away from OEC near lumen in water channels

### 4.1. Characterization of the channels away from the OEC

The network analysis shows two regions, one where the network is highly interconnected near the OEC followed by well separated changes moving to the lumen. Roughly the large channel separates near CD of D1-E329 10.6 Å from the OEC Ca; the narrow channel at Oγ of D1-S169, 6.4 Å from Mn4; the broad channel separates from the interconnected region near Cδ of D1-E65 13.6 Å from Mn4. The examples provided in Table 2 show water chains by which protons can exit to the lumen without amino acids serving as direct participants. However, several acidic residues can serve as intermediary proton loading sites. In addition, hydrogen bonds to non-Grotthuss competent sidechains help to organize the water orientations (Table 1).

### 4.2. Connection to the lumen via the narrow channel

The network analysis shows D1-S169 in the highly interconnected region that connects to D1-N338 within the separated narrow channel by a linear chain of ~6 water molecules. QM/MM calculations also found a well ordered water chain from O4 to D1-N338 [81]. Moving outward, D1-N338 and CP43-T335 are connected via 4 waters (Figure 2A). Thus, the narrow channel is indeed narrow. The narrow channel exits to the lumen near PsbU-E93, D96, CP43-K339 (Figure 3A).

On the surface, there are many hydrogen bond connections near the narrow channel exit. CP43-K339 and PsbU-D96 form a direct salt bridge. CP43-K339 is also at the center of a water mediated network connecting PsbU-E93 and CP43-T335 and anchored by D1-N338 and PsbU-N99. The water mediated network also connects PsbU-N99 with CP43-D321 and PsbU-E93 with PsbO-D102 (Figure S2). PsbO-D102 is near the broad channel exit.

### 4.3. Connection to the lumen via the broad channel

The broad channel separates from the region around the OEC close to the narrow channel, with D1-D61 associated with both channels. D1-E65, D2-E310 and E312 are at the boundary between the highly interconnected region near the OEC and the channel proper. Moving along the broad channel we find D1-S177, D2-N318, D2-T316, PsbO- R152, D102, D99. The broad channel exits to the lumen through PsbO-D158, K188, D224, D222 and H228. These residues extend over 10 Å and are connected via clusters of waters. FTIR difference spectra support many of these residues being part of a long range hydrogen bond network [72].

Previous studies have analyzed the multiple clusters of Asp and Glu residues on the surface of PsbO near the broad channel exit which can serve as proton loading sites [34]. These are connected either by direct hydrogen bonds or via water molecules [34]. For instance, D2-E310 is a center of a surface network connected to the acids PsbO-D222 and D224, and the bases H228 and K188. There is a direct hydrogen bond trapping a proton between two acids PsbO-D102 and E97 as noted earlier [34].

### 4.4. Connection to the lumen via the large channel

D1-E329 can be viewed as the boundary between the highly interconnected region near the OEC and the large channel, which runs through CP43 and PsbV. Although the large channel has been considered to be less polar, CP43- E413, T397, H398, PsbV-S39, E83, K79 are in the large channel water mediated network. The channel exits to the surface near the bases PsbV-K103, K104 and K134. PsbV-K103, which forms a salt bridge CP43-E83, is at the center of a network that includes residues CP43-T397, E394, and H398.

### 4.5. Interconnections between the channels at their exits

The large channel exit is to a well separated region of PsbU and PsbV. In contrast, the narrow and broad channel entrances are near each other and their exits are interconnected by residues in PsbO and PsbU in an extended surface network. These surface connections are flexible and vary between MD snapshots. For example, transient connections are made between PsbU-E93 (narrow channel) and PsbO-D102 (broad channel).

### 4.6. Continuity of channel from the OEC to the surface

The network analysis shows that rather than starting out as three independent water channels, the pathways are highly interconnected by many waters near the OEC (Figure 2B and Figure 2C). The MD trajectories allow for the mobility of the waters in these channels to be assessed. With the exception of W_X_, which is directly hydrogen bonded to O4, all other waters in the interconnected area around the OEC are found to leave and be replaced by other water molecules during the 100 ns trajectory. Thus, despite their individual mobility, this water-mediated hydrogen bond network is very stable overall.

The network analysis compared the persistence of hydrogen bonding connections in multiple MD snapshots in the separated channels moving towards the lumen. The broad channel forms a long, unbroken hydrogen bond pathway in 90% of the snapshots, while the narrow and large channels each become disconnected near the lumen in 30-40% of the snapshots. The break in the narrow channel occurs near CP43-K339 while for large channel, the hydrogen bond connection gets broken near PsbV-S39 and K103. In each case the connections are broken by transient dehydration events.

### 4.7. Proton affinity and protonation states of acidic and basic residues

MCCE keeps the protonation states of all amino acids in equilibrium with the pH and the hydrogen bond network. Most acidic and basic residues in the region around the OEC retain their expected protonation states. However, earlier MCCE studies identified some residues with different equilibrium ionization in the S1 state [27]. These are listed in the methods section and are fixed in the MD trajectory. All residues except for the primary amino acid ligands of the OEC are free to take any protonation state in the MCCE network analysis.

There are numerous acidic and basic residues within the highly interconnected region near the OEC (Table 1). Previous calculations of the proton affinity of residues in this region showed D1-D61, E65 and E329, which are all well connected to the network, all have significantly higher proton affinity than Asp or Glu in solution, although they are all predominantly anionic at pH 6 [10,27]. The higher proton affinity lowers the barrier for them to bind and release protons, facilitating proton transfer through the network entering the broad channel. Earlier continuum electrostatics based calculations [10] found the pK_a_s of D1-D61, E65, D2-E312, D2-K317, D59, R64, PsbO-R152 and D224 increase moving outward in the broad channel, again supporting their ability to function as proton loading sites.

### 4.8. Relative energy of hydronium in water channels

The three channels are all interconnected near the OEC (Figure 2B and Figure 2C) and proton transfer pathways can be identified to the lumen in each (Table 2). While the connections through the broad channel are less likely to be broken than those in the other two channels, the decision of which is used to transport protons may depend upon the relative energy of a positive charge in each channel. The free energy of replacing each water with a hydronium equilibrated with surrounding water molecules and amino acid residues by MC sampling was determined (Figure 1B). In each snapshot the energy of hydronium was obtained at 7-8 locations in each channel moving from the OEC to the lumen. Figure 5A shows the average energy of the hydronium for all the snapshots at different positions. Here the hydronium energy is binned into four 5-kcal groups from 0-20 kcal to provide a qualitative view of the energy of the energy at many locations. In a Grotthuss proton transfer mechanism each water in the chain is only transiently associated with having 3 protons so we do not wish to suggest a hydronium proper is moving through the channel. Rather these calculations should be viewed as probing the relative energy of a positive defect moving through the network. The excess proton can move through the water network or be found on an amino acid proton loading site [10].

**Figure 5.**
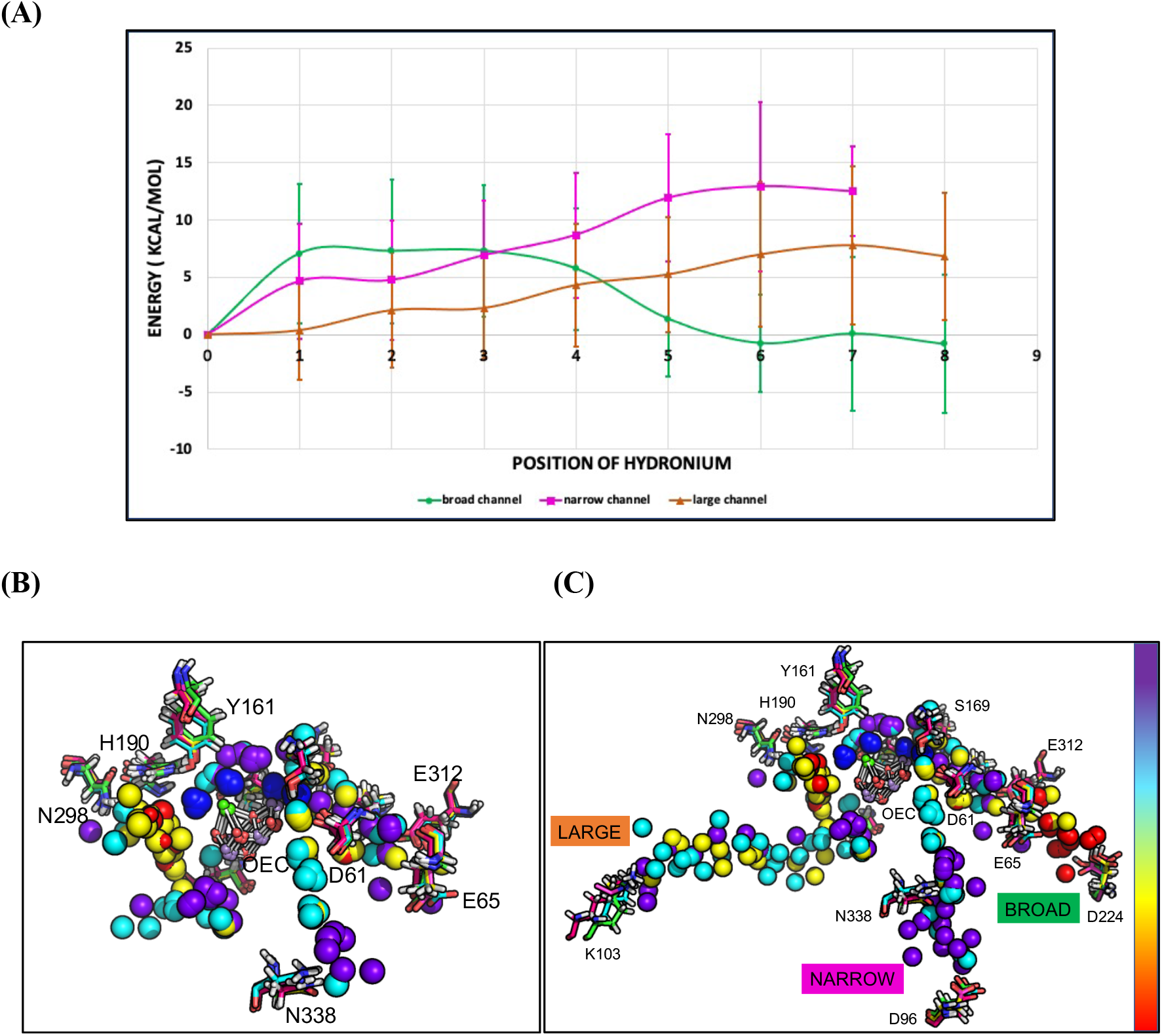
**(A)** Energy of PSII in MC sampling with hydronium at different positions moving into and along the broad, narrow and large channels. The average values from ten snapshots are shown with standard deviation shown as error bars. The reference energy (at position 0) is for the full protein with no hydronium plus the energy of an isolated hydronium in water at pH 8. The amino acids near each position are described in the text. Orange: Large channel; Magenta: narrow channel; Green: broad channel. **(B)** Energy landscape for water molecules near OEC (**left**) and reaching to the lumen (**right**) for all the waters in the channels. The results from 5 MD snapshots are superimposed. The water oxygens are colored to show the relative energy of the protein with hydronium with red<yellow<cyan<purple. Red is the most favorable and each color represents a 5 kcal/mol energy range. D1-D61, E65 and D2-E312, PsbO-D224 (broad channel), D1-S169, N338, PsbU-D96 (narrow channel) and D1-H190, Y161, N298 and PsbV-K103 (large channel) are shown. Figure S3 shows energy plots for individual snapshots.

Figure 5A shows the averaged energy at 8 locations from five superimposed snapshots, chosen by meta-analysis of the trajectory. The reference energy (position 0) is the sum of the energy of the protein (without hydronium) plus that of hydronium in solution. To differentiate the channels in the intertwined network near the OEC, the broad channel is considered to start near the waters of Mn4 moving towards PsbO-D224, while the narrow channel starts near D1-S169 moving towards CP43-K339 and PsbU-D96 and the large channel starts near Ca moving towards the PsbV-K103. Figure 5B colors the relative energy of each water in each superimposed snapshot. The proton affinity decreases from red to yellow to cyan to blue. Regions more favorable for hydronium (red and yellow) remain distinct from those that are less hospitable (cyan and blue).

Moving along the narrow channel, positions 1, 2 and 3 (yellow and cyan in Figure 5B) are near D1-S169 and D1-D61. While the less favorable positions 4 and 5 (cyan and purple) are close to D1-N338 and CP43-T335; unfavorable energy is found near positions 6 and 7 (cyan and purple) close to CP43-K339 and D1-D96. Dehydration events in the MD trajectory take place near position 6. Thus, although the region of the interconnected network close to the narrow channel is favorable for a positive charge, the channel itself is not.

In the broad channel, positions 1 and 2 (cyan and purple) are near to D1-V185 and are relatively unfavorable for hydronium compared to positions 3 and 4 (yellow and cyan) near D1-N181 and D61. Positions 5, 6, 7 near D1-E65 and D2-E312 (yellow and red) and 8 (red) near to PsbO-D224 are quite favorable for a positive charge. The increasingly favorable energy of a proton moving along the broad channel agrees with earlier calculations that found the protonation of acidic residues became more favorable moving outward along the broad channel [10]. The pK_a_ of the acidic amino acids and the free energy of the hydronium probe both show the channel is relatively hospitable to a positive charge.

Tracing hydronium through the large channel shows a favorable region at positions 1-3 (yellow and red) near the neutral D1-H190, N298, D1-D342, D1-Q165 and W4. However, moving outward the free energy of the hydronium probe increases. Positions 4, 5 and 6 are near D1-E329 and CP43-E413 (yellow, cyan and purple) while 7 and 8 are near the channel exit PsbV-K103 (cyan and purple). The dehydration of the large channel occurs near position 7.

The relative energy for the initial positions of hydronium near to the OEC in all the water channels are within the error bars of the distribution of energies of nearby hydronium probes. Favorable positions are W_X_ near to D1-S169 in the narrow channel, at waters near D1-N298 in the large channel and waters near D1-D61 in the broad channel (Figure 5B). However, as the protons move away from the OEC, the energy is more favorable in the broad channel while it continually increases moving through the narrow and large channels. This means the that broad channel has a surmountable barrier to proton transfer while proton transfer in the narrow and broad channels is entirely uphill.

### 4.9. Comparison of network analysis results with network proposed by FTIR

FTIR spectra are very sensitive to protonation states and hydrogen bond strengths in networks of waters and amino acids. As a raw FTIR spectrum is extremely congested, the difference between spectra in different S-states, and double difference spectra comparing wild-type protein with that labeled with different isotopes or modified by site-directed mutations are used to probe the connectivity of the network that responds to the OEC reactions. Many mutations have been made in the region of amino acids separated by as much at 20 Å around the OEC shown in Figure 6. These include: the large channel residues D1-Q165E [80], D1-E329Q [72], N298A [48,74], narrow channel residues D1-N87A [69], D1-S169A [68] and broad channel residues D1-D61A [70], E65A [72], N181A [75,77] and D2-K317A, K317R, K317Q, K317E [71]. It was found that mutations of the residues in the inner, interconnected network described here produce similar changes in OEC function as well as in the FTIR signatures of the hydrogen bond networks [72,75].

**Figure 6.**
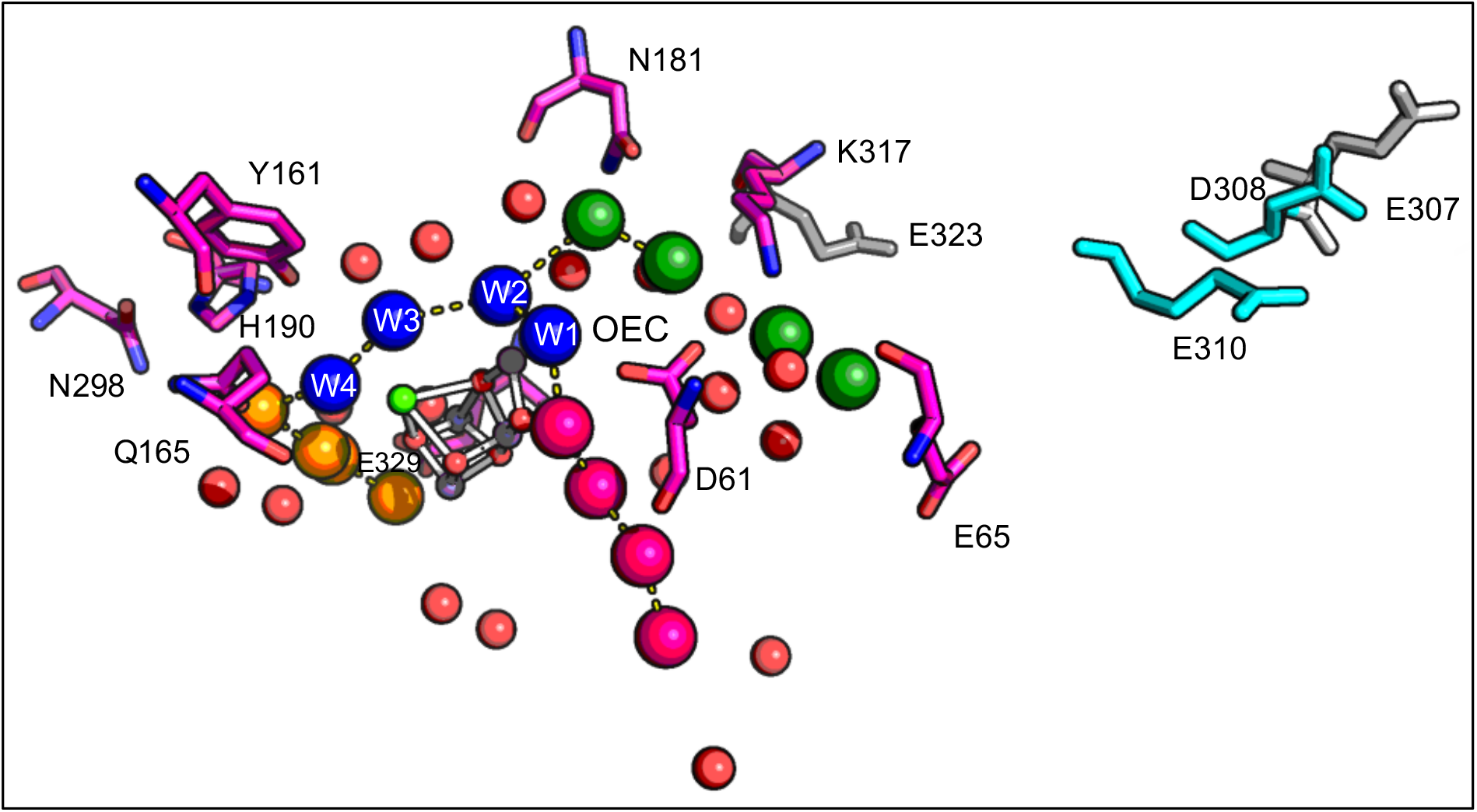
Residues identified as being members of a long-range hydrogen bond network. The residues are: D1-D61 [70], E65 [72], N181 [75,77], D2-K317 [71] (broad channel): D1-N87 [69], S169 [68] (narrow channel): D1-Y161 [82], H190, Q165 [80], N298 [74], E329 [72] (large channel). Residues and waters are colored for the channel with which they are most closely associated; Blue: W1, W2, W3 and W4; Red: water molecules not assigned to a specific channel. Cyan and Grey: residues whose mutation does not alter the FTIR difference spectra. Cyan: D2-E310, D308 are connected to the end of the broad channel in the MCCE derive network; Grey: D2-E307 and E323 are not in MCCE network.

Mutations of D2-E307; D2-D308; D2-E310 and D2-E323 do not modify the FTIR spectrum [80]. D2-E310 and D2-D308 are in the MCCE network, but the former is near the end of the broad channel and the later on the surface. D2-E307 and E323 are not connected to the network found here, nor do they affect the FTIR difference spectra (highlighted in grey color) (Table 1). Note that D2-E323 is on the surface and is farther away from D2-K317.

## 5. Conclusions

The results provide characterization of the proton egress pathway by combined classical MD, MCCE and network analysis. An unexpected conclusion is that the three channels are highly interconnected in a region reaching 10± 2 Å from the edge of the OEC (Figure 2B). In addition, the free energy of a hydronium probe replacing all of the waters around the OEC is similar at most sites in this region, independent of which is the nearest channel entry. The three channels do disentangle themselves as they move towards the lumen. However, the entries to the narrow and broad channels are close together near the OEC, connected by D61 and the are also interconnected by surface hydrogen bond networks at their exits.

The results reported here support the broad channel as the preferred proton exit. It retains a lower, energy for hydronium to the end of the channel (Figure 5B). This is in agreement with earlier continuum electrostatics studies [10] that used the pK_a_’s of amino acids as a probe, showing increasing proton affinity moving along the broad channel to the lumen. In contrast, the energy of the hydronium probe in the narrow and large channels increases moving towards the exit. Additionally, the broad channel water chains are rarely broken in the MD trajectory, while the narrow and large channels become transiently disconnected by dehydration events. This result supports earlier MD simulations that found a long-range hydrogen bond network starting from Mn4 of the OEC extending to the PsbO subunit present in the lumen through the broad channel [10,12,34].

## Acknowledgements

We would like to acknowledge Dongyue Liu from Boston University for insightful discussions and assistance in the set-up of the PSII MD calculations in CHARMM and OpenMM. The authors would like to acknowledge financial support from the Division of Chemical Sciences, Geosciences, and Biosciences, Office of Basic Energy Sciences, U.S. Department of Energy, Photosynthetic Systems. Experimental work was funded by DESC0001423 (M.R.G. and V.S.B) and DE-FG02-05ER15646 (G.W.B.). V.S.B. acknowledges DOE high-performance computing time from NESRC. The MD simulations on monomeric PSII used the computing time from Memorial Sloan Kettering Cluster and resources at Oak Ridge National Laboratory, supported by the Office of Science at DOE under the contract no. DE-AC05-00OR22725, made available via the INCITE program.

## Supporting Information

S1 is a figure of PSII highlighting the portion of protein used for MCCE calculations, S2: Hydrogen bond network for 9 snapshots mediated by 4W, S3: Energetic profile for individual snapshots 1-10.

