## Supplementary material for "Proton egress pathway during the S_1_–S_2_ transition of the Oxygen Evolving Complex of Photosystem II": Supplementary_information.pdf

**S1.** Figure of PSII highlighting the portion of protein used for MCCE calculations

**S2.** Hydrogen bond network for 9 snapshots mediated by 4 water molecules

**S3.** Energetic profile for individual snapshots 1-10

#### S1. Figure of PSII highlighting the portion of protein used for MCCE calculations

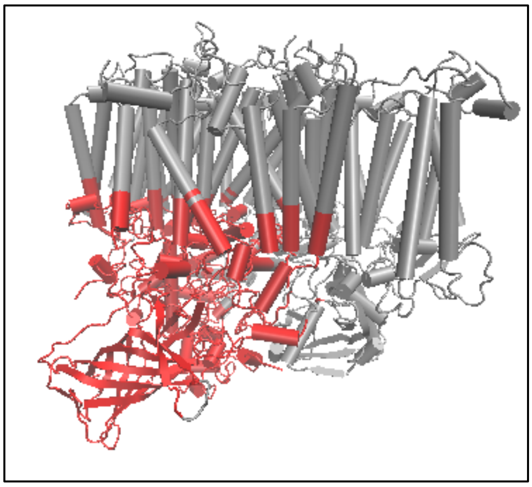

**Figure S1** Structure of PSII (PDB: 4UB6) [1] in gray color and the highlighted red portion of size 88 Å × 70 Å × 65 Å of protein used for MCCE calculations.

#### S2. Hydrogen bond network for 9 snapshots mediated by 4 water molecules

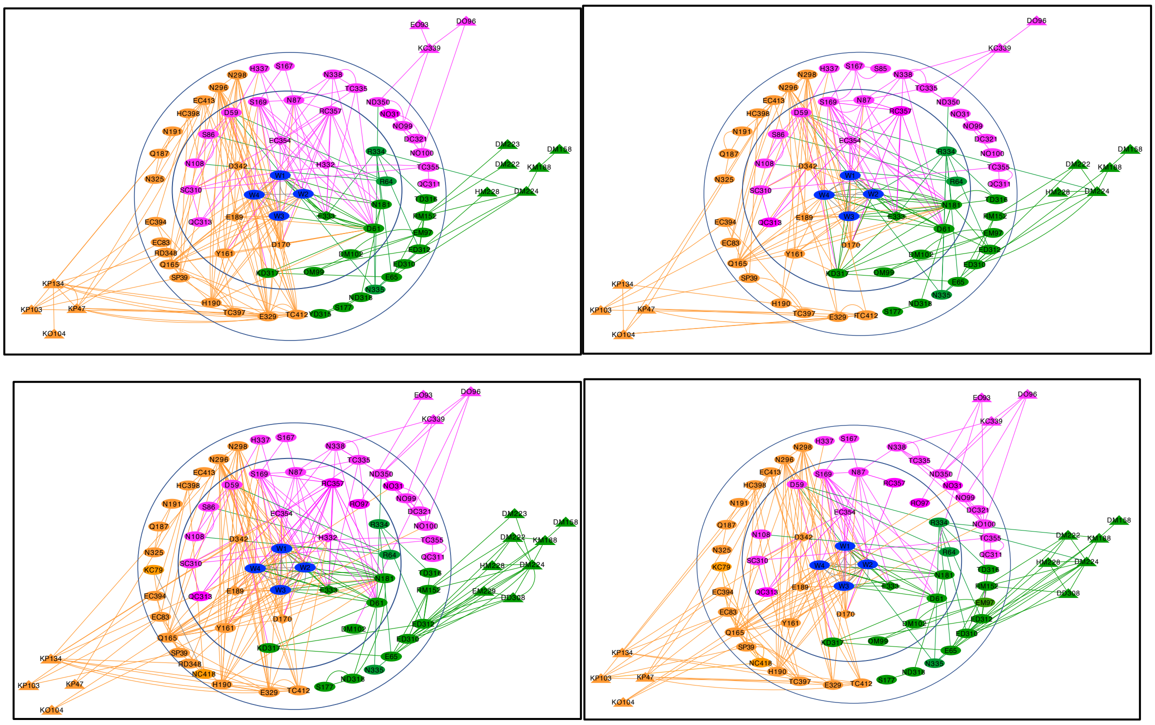

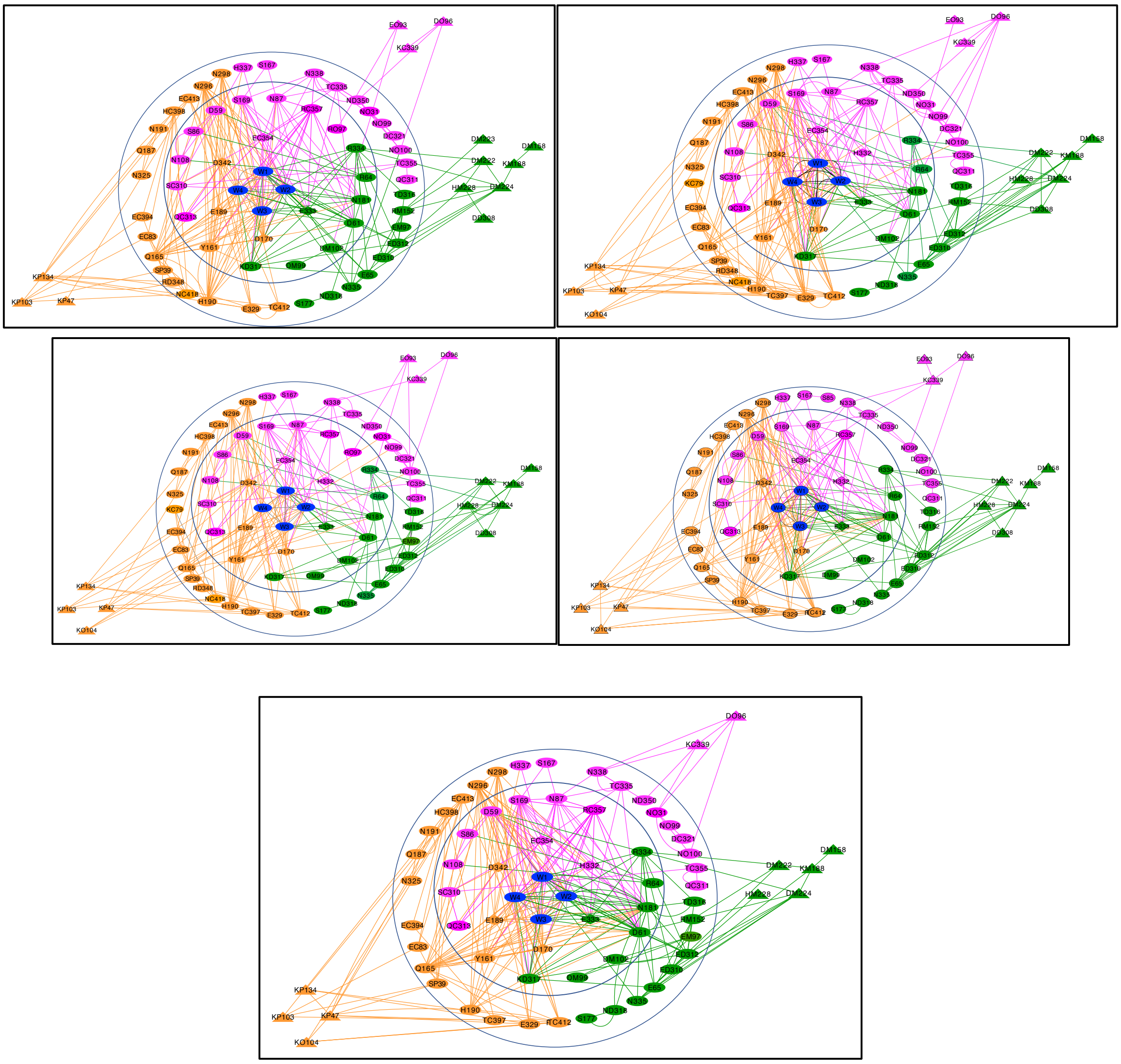

**Figure S2** Network of hydrogen bond connections from the OEC to the lumen found in MCCE calculations for 9 MD snapshots. Nodes are labeled as Residue type Chain designation Residue number. No chain designation indicates D1. Chain C is CP43; D is D2; M is PsbO; and P is PsbV, O is PsbU. For example, EC354 is CP43-Glu354. Diamonds are primary ligands and triangles are residues with at least 20% of their surface exposed to the lumen. Lines show hydrogen bond connection mediated by 0-4 waters. The inner circle encloses highly interconnected residues near the OEC. Connections between residues nominally in different channels are seen. The outer circle encloses residues in their separated channels. Beyond the outer circle are residues connecting the channel and the surface.

### S3. Energetic profile for individual snapshots 1-10

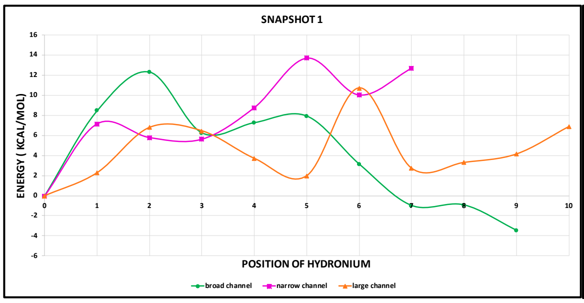


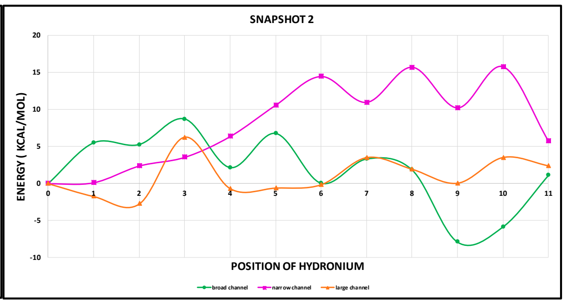


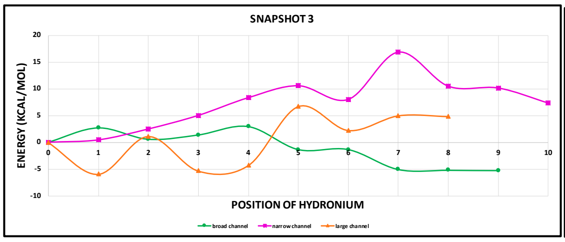


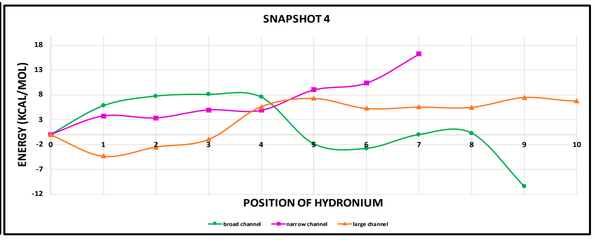


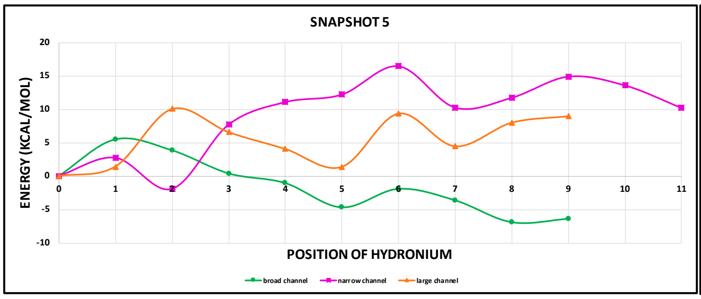


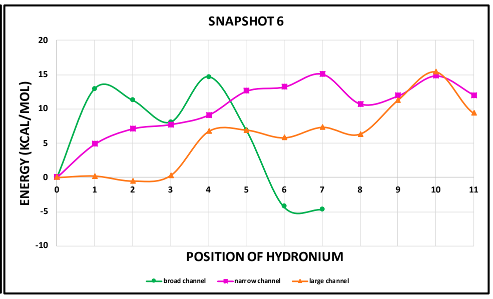


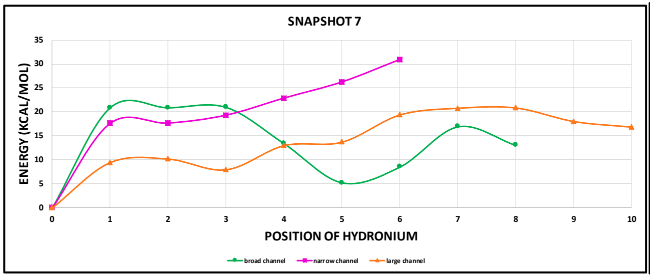


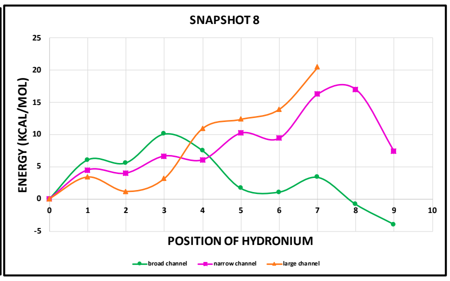


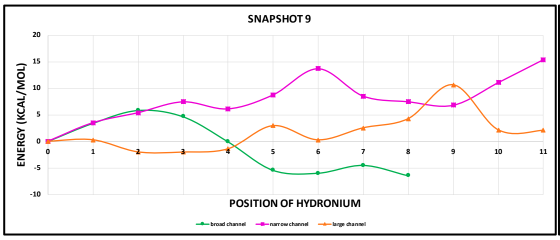


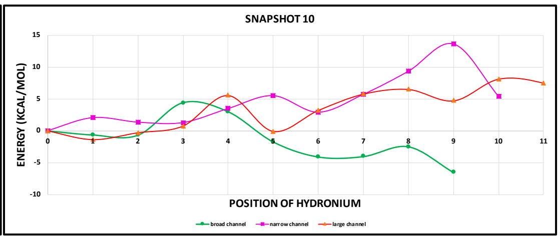

**Figure S3** Free energy profile for hydronium at different positions in the broad, narrow and large channels for ten individual snapshots. Large channel is orange, narrow channel in magenta color while broad channel is in green. x-axis is the position of hydronium at various positions in all three water channels moving away from the OEC. y-axis is the energy (in kcal/mol) of hydronium. The reference energy (at position 0) is for the full protein with no hydronium plus the energy of an isolated hydronium in water.
